# Invasive species undermine the bioark hypothesis in a low-latitude urban biodiversity hotspot

**DOI:** 10.64898/2026.05.24.727550

**Authors:** Luke Edenborough, John Paul Hellenbrand, Samantha Kennett, Sofia Cuenca Rojas, Clint A. Penick

**Author notes:** Correspondence to: Luke Edenborough, 301 Funchess Hall, Auburn, AL 36849.

## Abstract

Urbanization often reduces local biodiversity, yet cities can maintain surprisingly high total species richness at broader spatial scales. This paradox has led to the “cities-as-bioarks” hypothesis, which proposes that protected remnant habitats within cities can function as refuges for native species. However, most evidence supporting this hypothesis comes from higher latitudes, and it remains unclear whether remnant habitats in low-latitude cities can serve the same role. We surveyed ant communities across 60 sites in Atlanta, Georgia (USA), spanning streetscapes, manicured parks, and forested parks, to test whether relatively undisturbed urban forests support the highest native diversity. Contrary to the bioark prediction, native species richness was lowest in forested parks and highest in habitats with intermediate disturbance. Community composition varied with habitat structure, but surface temperature was not a significant predictor of richness or composition. Instead, native abundance and richness declined strongly with increasing abundance of the invasive Asian needle ant, *Brachyponera chinensis*, a forest-adapted invader capable of dominating relatively intact habitats. In contrast, two other invasive species, the red imported fire ant *Solenopsis invicta* and the Argentine ant *Linepithema humile*, were largely restricted to more disturbed habitats and had comparatively weaker associations with native diversity loss. These findings refine the bioark hypothesis by demonstrating that habitat protection alone may be insufficient to conserve insect diversity in warmer regions, and instead must be paired with active invasive species management.

## Introduction

Although biodiversity frequently declines with increasing urban intensity (Sanllorente *et al*., 2025), studies of urban insects show that total species richness at the city scale can remain high (Baldock *et al*., 2015; Savage *et al*., 2015). This paradox arises, in part, because cities are mosaics of habitats that differ in urban intensification and management (Sattler *et al*., 2011). Protected natural habitats within city bounds may function as “bioarks” that preserve native species within otherwise heavily altered landscapes (Brassard *et al*., 2021; Diamond *et al*., 2023). Most support for the cities-as-bioarks hypothesis comes from northern latitudes. Because most of the world’s biodiversity is concentrated at low latitudes, testing the bioark hypothesis in these regions is essential for understanding how urbanization reshapes global diversity. Low-latitude areas may experience stronger pressures from urban heat and invasive species, potentially limiting their capacity to sustain native communities (Guo *et al*., 2021; Rajesh *et al*., 2022; Youngsteadt *et al*., 2016). Whether remnant natural habitats in low-latitude cities can function as bioarks under these combined pressures remains unresolved.

Urban climate represents one major pathway through which cities restructure insect communities. Urbanization elevates temperatures and alters microclimatic conditions, often increasing thermal stress in exposed habitats (Sears *et al*., 2007). However, the ecological consequences of urban warming may vary with latitude, altering the strength or direction of these effects. For example, in a global analysis of bird and plant diversity, Aronson *et al*., (2014) found that the magnitude of urban biodiversity loss depended strongly on regional climate and biogeography, and Leveau *et al*. (2017) demonstrated that urbanization flattened latitudinal patterns of avian species richness across the southern Neotropics, with the largest urban-rural differences occurring at lower latitudes. Similar patterns have emerged in studies of insects and other arthropods, for example, Youngsteadt *et al*. (2016) found that urban warming increased insect abundances at high latitudes but had increasingly negative effects with decreasing latitude. In a study focused on ants, Penick *et al*. (2025) observed consistent declines in species richness with urban heat across cities, but there were potentially greater absolute losses in species-rich, low-latitude regions. These findings suggest that urban heat may relax thermal constraints in cooler cities while intensifying them in already warm regions where species operate closer to their physiological limits. Stronger negative effects of urban warming at low latitudes may therefore undermine the capacity of cities to function as bioarks, especially if warming extends into remnant habitats.

Biological invasion represents a second major pathway through which cities restructure insect communities. Invasive species are major drivers of biodiversity loss worldwide and impose substantial economic costs (Angulo *et al*., 2022; Gruber *et al*., 2021; Senator & Rozenberg, 2017). Urban ecosystems, in particular, tend to support high densities of invasive ants (Gochnour *et al*., 2019), which thrive in disturbed habitats and can suppress native species through competition, predation, and resource monopolisation (Carpintero & Reyes-López, 2008; Holway *et al*., 2002). In a large-scale analysis of urban ant diversity, Perez *et al*. (2022) found that urbanization had stronger negative effects on diversity at low latitudes. Although that study did not directly test the role of invasive ants, it documented greater divergence between urban and non-urban communities in low-latitude cities, suggesting a shift toward communities dominated by non-native species. These patterns suggest that low-latitude cities may experience stronger invasion pressure than their high-latitude counterparts, which may erode the bioark potential of low-latitude cities.

Here, we surveyed ant communities in Atlanta, Georgia (USA), to test whether protected remnant habitats can sustain high native insect diversity (the bioark hypothesis) or whether this potential declines at lower latitudes. Atlanta lies within the largest urban corridor in the southeastern United States (Terando *et al*., 2014) and hosts three prominent invasive ants: red imported fire ants (*Solenopsis invicta* Buren 1972), Argentine ants (*Linepithema humile* (Mayr, 1868)), and the more recently established Asian needle ant (*Brachyponera chinensis* (Emery, 1895)). Of these, *B. chinensis* is unique in its ability to colonise and dominate intact forest habitats, making it a potential driver of diversity loss even in low-disturbance environments (Guénard *et al*., 2018; Guénard & Dunn, 2010; Kanes *et al*., 2025; Warren *et al*., 2015). Under the bioark hypothesis, we predicted that forested parks would support the highest native diversity. If urban climate effects dominate, we expected steeper declines in diversity with increasing urban heat. If invasion exerts stronger control, we predicted reduced native diversity in sites where invasive species abundance is high. Together, these predictions allow us to evaluate whether urban habitats in a low-latitude city function as bioarks and how their responses to climate and invasion differ from those observed at higher latitudes.

## Methods

### Study sites

We sampled ant communities at 60 sites in Atlanta, Georgia (USA) during June-August 2021 and June 2023 (**Table S1**; **Fig. 1**). We selected sites to represent three common urban habitat types that vary in disturbance: streetscapes (high disturbance), manicured parks (moderate disturbance), and forested parks (low disturbance). We defined streetscapes as highly fragmented areas under 5,000 m², such as traffic islands, roadside plantings, and greenspaces near buildings. We defined manicured parks as green spaces over 5,000 m² with mixed canopy cover and an understory dominated by mowed grass or planted woody vegetation. Finally, forested parks were forested areas over 5,000 m² categorised by minimal maintenance, dense canopy cover, and the ground being covered primarily in leaf litter. The dominant plants in forested parks were white oak (*Quercus alba* L.), loblolly pine (*Pinus taeda* L.), American beech (*Fagus grandifolia* Ehrh.), poison ivy (*Toxicodendron radicans* [L.] Kuntze), English ivy *(Hedera helix* L.), and Virginia creeper (*Parthenocissus quinquefolia* [L.] Planch.). In total, we sampled 18 streetscapes, 20 manicured parks, and 22 forested park sites across the city across all sampling methods.

**Fig. 1.**
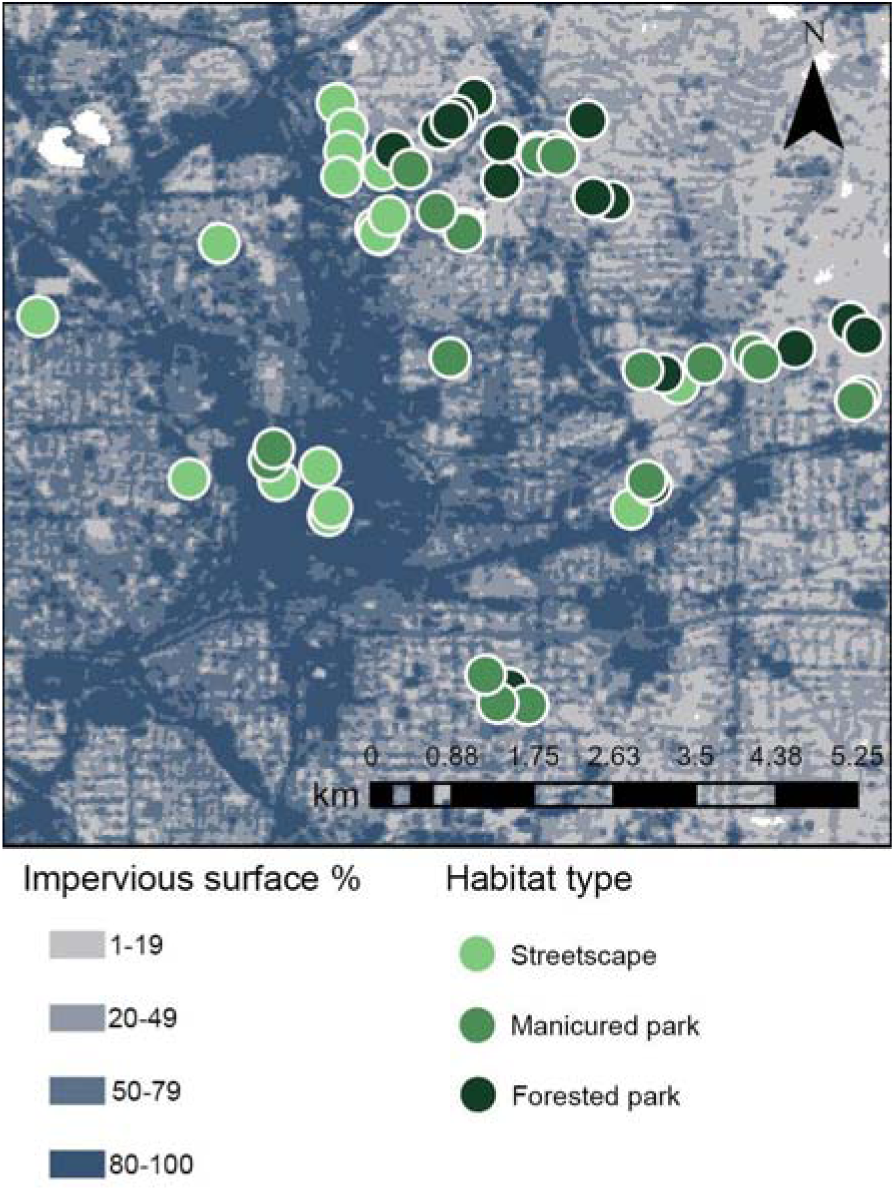
Spatial distribution of impervious surface cover across Atlanta, Georgia (USA), to a resolution of 30 metres using Dewitz (2023). Darker shading indicates higher percentages of impervious surface. Sampling sites are shown and color-coded by habitat type.

To quantify environmental variation among sites, we ran two perpendicular transects (20 m each) that intersected at the centre of each site. At 5-meter intervals along each transect, we measured leaf litter depth and ground cover type by visual inspection (categories included: bare ground, mulch, leaf litter, fine vegetation, or coarse vegetation). Following methods from Penick *et al*., (2025), we computed a vegetation complexity index (VCI) as the Shannon diversity index (*H*) of vegetation categories, where each vegetation type was treated as a ‘species’ and it’s abundance was the percentage of cover. At the centre of each site, we also recorded canopy cover using the ‘Percentage Cover’ iOS app (Mignanelli, 2021) at each cardinal direction, taking the average. We supplemented on-site data with NLCD and Landsat Collection in ArcGIS Pro to extract percent impervious surface and surface temperature with a resolution of 30 m (Dewitz, 2023; U.S. Geological Survey, 2021). Surface temperature values were extracted from all available cloud-free Landsat scenes collected between June 1 and August 31, 2023.

### Ant Sampling

To capture the widest diversity of ant species across habitats, we used a combination of standard ant sampling methods (Gotelli *et al*., 2011; **Fig. S1**) across two sampling periods. During June-August 2021, we conducted timed hand collection, baiting, and leaf litter extraction at all sites (**Table S1**). In June 2023, we conducted pitfall trapping at most previously sampled sites and included twelve additional sites not sampled in 2021 (**Table S1**).

For hand collection, we used forceps and aspirators to collect ants across microhabitats (e.g., under rocks, logs, within leaf litter, and up tree trunks) within our 20 x 20 m plots for a cumulative total of one hour per site (sum of time each individual spent sampling). Specimens were preserved in 70% ethanol for later identification. For baiting, we provided a multi-bait buffet that included olive oil, 20% L-glutamine, 20% sucrose, 5% NaCl, and tap water. We arranged the baits in a circular pattern spaced evenly apart (10 cm) at the center of our two transects and recorded the species and number of individuals feeding at each bait after one hour (**Fig. S1c**). We retained voucher specimens for any ants that could not be identified in the field. For leaf litter sampling, we collected a 1 m² quadrat from the area with the densest visible litter. We sifted the litter and placed it in a Winkler extractor (Ivanov et al. 2010) for 48 hours to collect ants and other arthropods into propylene glycol. In 2023, we deployed pitfall traps along a 10-meter transect at each site where ground conditions allowed. Plastic specimen cups (5 cm diameter) were filled halfway with water mixed with dish detergent to break surface tension, and then placed flush with the soil surface at 5 m intervals. The traps were retrieved after 48 hours, the contents were strained, and all remaining material was stored in 70% ethanol for later sorting and identification.

We identified all ants to species using the Mississippi Entomological Museum’s dichotomous key to the ants of the southeastern United States (MacGown, 2024). Voucher specimens were deposited in the Auburn University Museum of Natural History.

### Statistical Analyses

All analyses were performed in R 4.4.1 (R Core Team, 2025). To assess differences in species richness among habitat types, we computed sample-based species accumulation curves (random site accumulation, 999 permutations) using the ’vegan’ package (Oksanen *et al*., 2025) based on pooled species counts from baiting, leaf litter extraction, and hand collection (pitfall traps were excluded as not all sites were sampled using this method). We also compared site-level richness among habitat types using the same sampling methods with a Kruskal-Wallis rank-sum test. To compare ant abundance among habitats, we calculated mean per-site pitfall abundance (averaged across all species records at each site) and fit a Poisson generalized linear model (GLM) with a log link function, and differences among habitats were followed up with estimated marginal means (EMMs) and pairwise contrasts with Benjamini-Hochberg adjustment.

To evaluate the effects of urban heat and other environmental factors on community composition and species richness, we used a Jaccard distance-based redundancy analysis (db-RDA). Predictor variables were retained only after confirming acceptable collinearity (variance inflation factors < 10). Overall and marginal significance were assessed using ANOVA-like permutation tests (999 permutations). For variables with acceptable collinearity, we also tested bivariate relationships between site-level richness and six environmental predictors (percent canopy cover, percent impervious surface, percent leaf litter cover, leaf litter depth [cm], surface temperature [°C], and vegetation complexity) using separate linear models (LMs).

To test the effects of invasive species on community composition as well as native ant abundance and species richness, we conducted a Jaccard db-RDA with site-level abundances of three invasive ants as predictors (*Brachyponera chinensis*, *Solenopsis invicta*, and *Linepithema humile*). Because abundance estimates were derived from pitfall traps, these analyses were restricted to pitfall samples. Significance of overall and marginal effects was assessed using ANOVA-like permutation tests (999 permutations). We followed significant effects with Poisson GLMs (log link) to test the effects of invasive species abundance on native ant abundance and native species richness with habitat type included as an additional predictor.

## Results

### Community richness and abundance across habitats

We identified 54 ant species from 22 genera across all sampling methods and sites (**Table S2**). Contrary to expectations, species richness did not peak in the least disturbed habitat (forested parks) but instead was highest in habitats with intermediate urban stress (manicured parks; **Fig. 2a**); richness was 32 species in streetscapes, 33 in forested parks, and 40 in manicured parks. Mean site-level richness was highest in manicured parks, intermediate in streetscapes, and lowest in forested parks (**Fig. 2b**), though differences were not significant (Kruskal-Wallis, *N =* 47, *df =* 2, χ*^2^ =* 5.35, *p =* 0.069). Ant abundance varied significantly among habitats based on pitfall samples (Poisson GLM, *N =* 34, *df* = 31, residual deviance = 124.25, *p* < 0.0001). Post hoc comparisons of estimated marginal means (EMMs) revealed that streetscapes supported higher mean per-site ant abundance than manicured parks (ratio = 2.22, SE = 0.37, *z* = 4.81, *p* < 0.0001) and forested parks (ratio = 1.70, SE = 0.27, *z* = 3.35, *p* = 0.001), while forested and manicured parks did not differ significantly (ratio = 1.31, *z* = 1.66, *p* = 0.097).

**Fig. 2.**
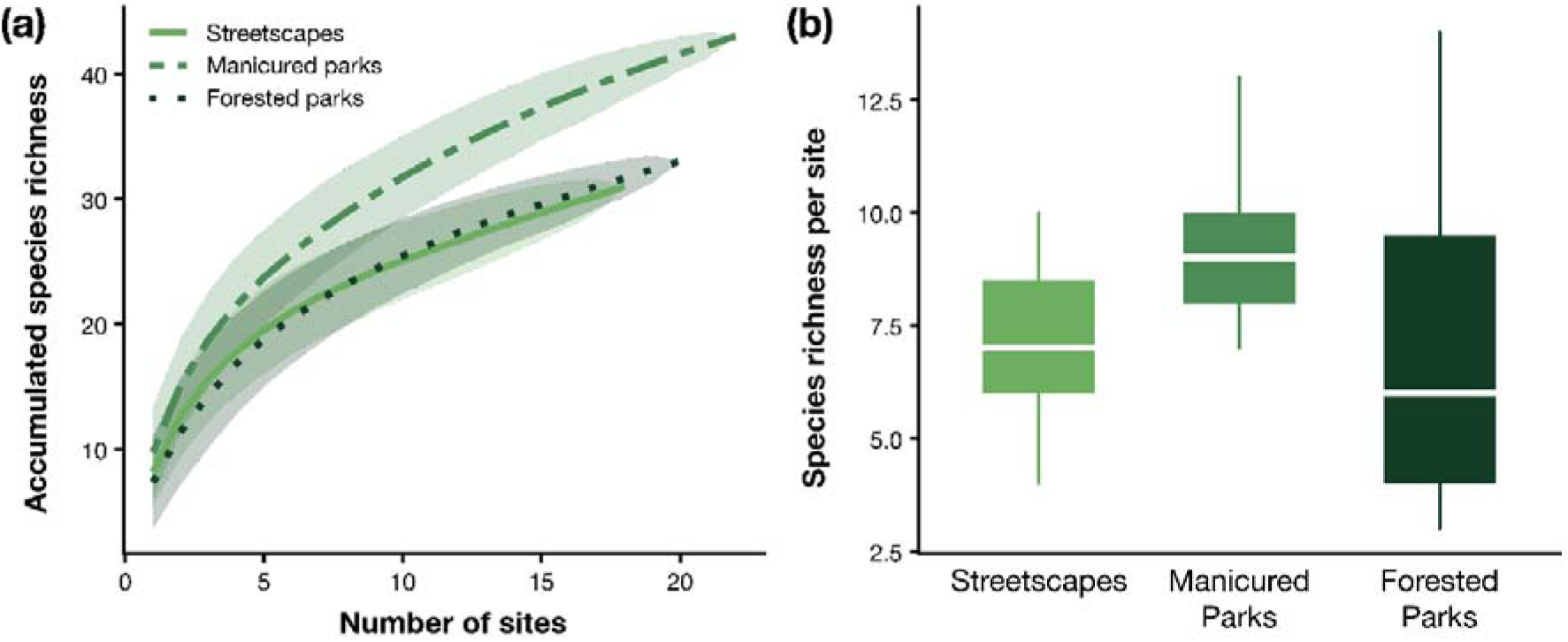
Ant species richness across habitat types. **(a)** Species accumulation curves comparing ant species richness among habitat types. Shaded areas represent 95% confidence intervals. **(b)** Boxplots showing site-level species richness across habitat types (Median and interquartile range), though differences among habitat types were not statistically significant (Kruskal-Wallis, *p* = 0.069).

**Fig. 3.**
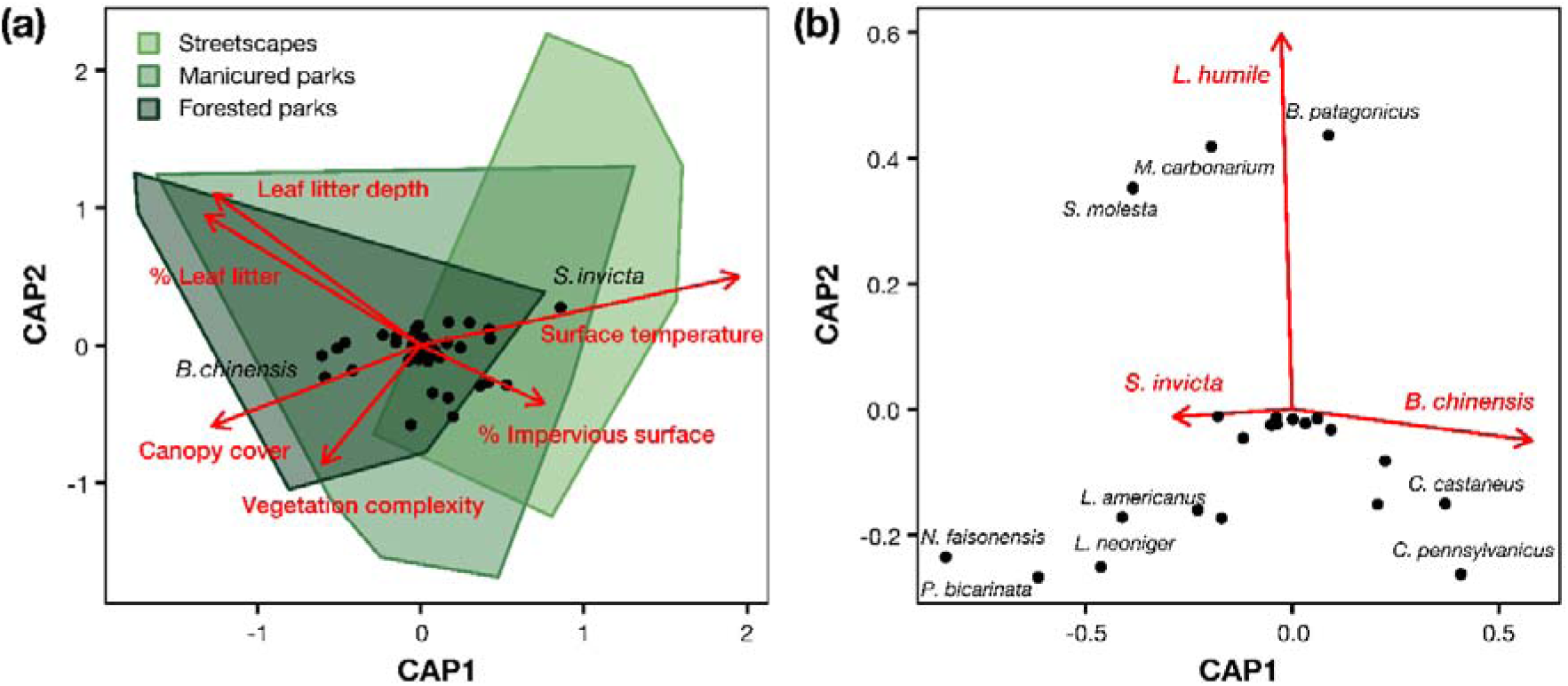
Abiotic and biotic drivers of ant community composition. **(a)** Jaccard distance-based redundancy analysis (db-RDA) of species composition constrained by environmental predictors; convex hulls indicate habitat types. **(b)** Jaccard db-RDA of pitfall-trap communities constrained by abundances of *Brachyponera chinensis*, *Solenopsis invicta*, and *Linepithema humile*. CAP1 and CAP2 are the first and second constrained axes of the db-RDA, which are the linear combinations of predictor variables that capture the largest fractions of variation in ant community composition. Predictors are shown in red.

**Fig. 4.**
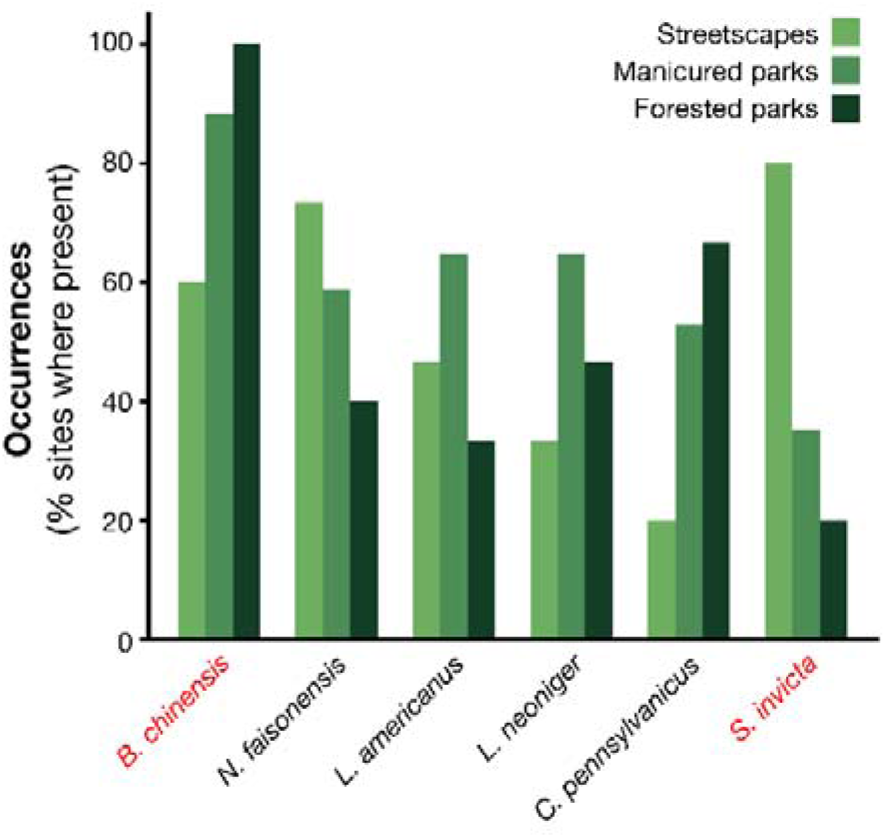
Occurrences of the six most frequently detected ant species in Atlanta by habitat type, ordered from most to least common (left to right). Invasive species are indicated in red text.

**Fig. 5.**
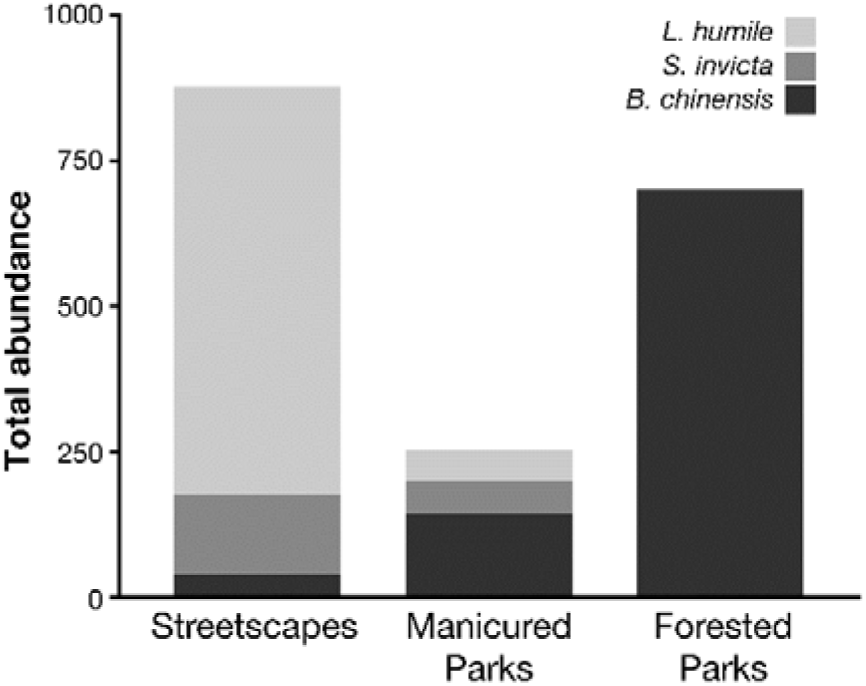
Stacked bar plot showing the mean abundance of the three main invasive ant species (*Brachyponera chinensis*, *Solenopsis invicta*, and *Linepithema humile*), grouped by habitat type (streetscapes, manicured parks, and forested parks).

### Abiotic drivers: urban heat and environmental variables

Ant community composition was significantly related to the environmental predictor set in a distance-based redundancy analysis (db-RDA) on Jaccard dissimilarities (ANOVA-like permutation test, pseudo-*F* = 2.03, *p* = 0.001; 999 permutations). Marginal permutation tests indicated that surface temperature (pseudo-*F* = 1.66, *p* = 0.039) and canopy cover (pseudo-*F* = 1.61, *p* = 0.042) were significant predictors of community composition, whereas leaf litter cover (*F* = 1.53, *p* = 0.061), leaf litter depth (*F* = 1.37, *p* = 0.094), and vegetation complexity (*F* = 1.39, *p* = 0.099) were marginally non-significant, and habitat category had no significant effect (*F* = 0.86, *p* = 0.739). Separate LMs of site-level species richness on each abiotic predictor showed that richness decreased significantly with increasing leaf litter cover (β = −0.050, *t* = −2.93, *p* = 0.005, *R*² = 0.16) and leaf litter depth (β = −2.21, *t* = −3.46, *p* = 0.001, *R*² = 0.21), but was not significantly related to surface temperature (*t* = 0.31, *p* = 0.761), canopy cover (*t* = −1.01, *p* = 0.316), impervious surface (*t* = 0.51, *p* = 0.611), or vegetation complexity (*t* = 0.41, *p* = 0.683; **Fig. S2**).

### Biotic drivers: invasive species

Ant community composition varied significantly with invasive ant abundance in a db-RDA constrained by abundances of *Brachyponera chinensis, Solenopsis invicta,* and *Linepithema humile* (ANOVA-like permutation test: *F =* 1.98, *p =* 0.001, 999 permutations; *R²* = 0.165, adj. *R²* = 0.082). Marginal tests indicated significant effects of *B. chinensis* (*F =* 2.99, *p* = 0.001) and *L. humile* (*F =* 1.50, *p =* 0.017), while *S. invicta* had no significant effect (*p >* 0.0552). Results from GLMs showed that native ant abundance declined significantly with increasing *B. chinensis* abundance (Poisson GLM, β = −0.0119 ± 0.0010 SE, *z* = −11.48, *p* < 0.0001; **Fig. 6a**). On the response scale, each additional *B. chinensis* individual corresponded to an ∼1.2% decrease in expected native abundance (exp(β) = 0.988), with an estimated baseline of ∼50 native individuals at sites where *B. chinensis* was absent (exp(intercept) = 49.9). The negative relationship between *B. chinensis* abundance and native abundance remained significant after including habitat as a covariate, although the effect size was reduced (Poisson GLM: β = −0.0042 ± 0.00126 SE, *z* = −3.36, *p* = 0.00078; **Fig. 6a**). After controlling for habitat type, *B. chinensis* abundance showed a marginal association with native richness (Poisson GLM: β = -0.0064 ± 0.0034 SE, *z* = -1.88, *p* = 0.0607; **Fig. 6b**).

**Fig. 6.**
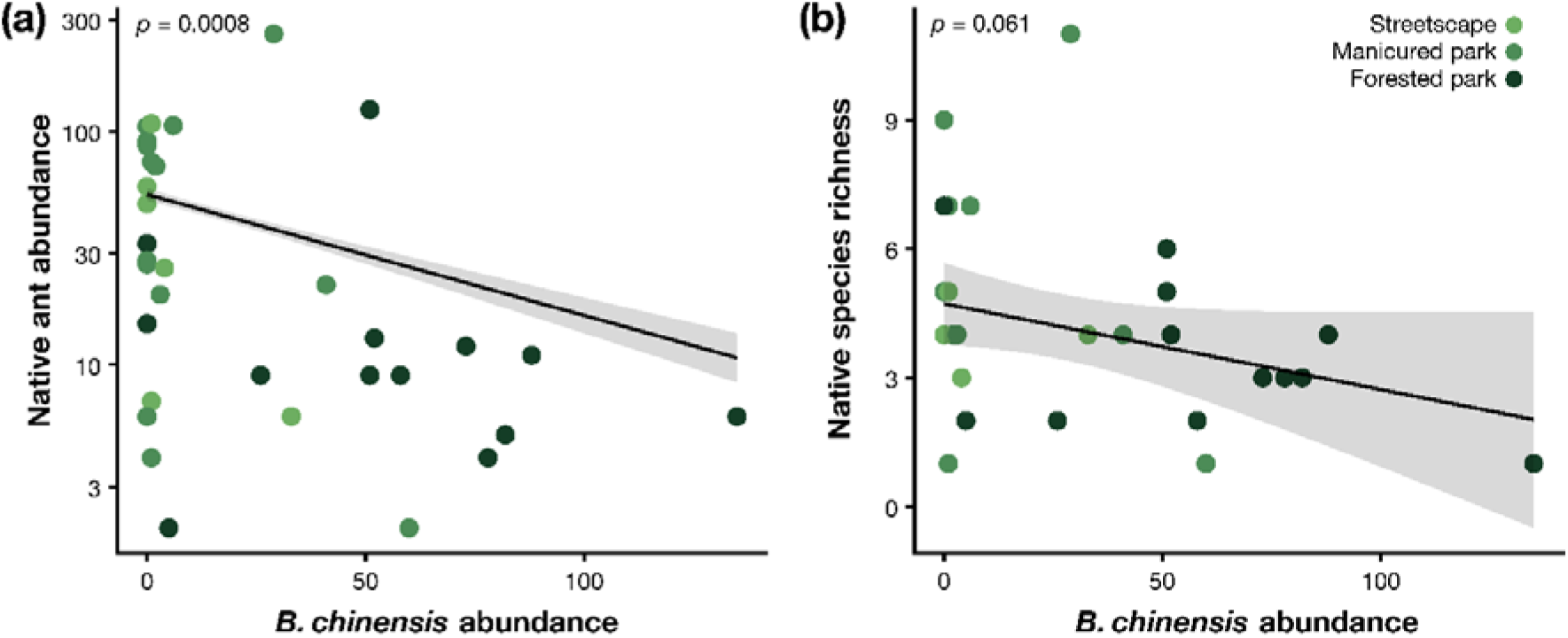
Relationships between *Brachyponera chinensis* abundance and **(a)** native ant abundance and **(b)** native ant richness. Plots show best fit line with 95% confidence intervals, and habitat type is indicated by colour. P-values are taken from Poisson GLMs with habitat type included as a predictor.

## Discussion

Although previous studies suggest that cities function as bioarks for insect diversity, our results indicate that this capacity may be reduced due to invasion. Consistent with the global latitude-diversity gradient (Griffith *et al*., 2025; Penick *et al*., 2025; Perez *et al*., 2022; Wallace, 1878), overall ant species richness was higher in Atlanta (55 species) than in comparably sampled northern cities, such as New York City, New York (USA), (40 species; Penick, 2021). However, within Atlanta, species richness was lowest in relatively protected forested parks. This pattern contrasts with findings from northern cities, where intact forest habitats typically support the highest native richness (Fenoglio *et al*., 2020; Sanllorente *et al*., 2025; Sattler *et al*., 2011; Savage *et al*., 2015). This decline in species richness was not related to common abiotic urban stressors, such as urban heat or impervious surface cover. Instead, native diversity and abundance declined with increasing abundance of the invasive Asian needle ant (*Brachyponera chinensis)*. This result aligns with broader patterns showing greater divergence between urban and non-urban ant communities at lower latitudes (Perez *et al*., 2022) and suggests that biological invasion may limit the ability of remnant habitats to function as bioarks in low-latitude cities, even where regional species pools are large.

Urban heat is widely recognized as one of the most consistent stressors in urban ecosystems (Cabon *et al*., 2024; Dietzel *et al*., 2025; Fenoglio *et al*., 2020; Penick *et al*., 2025; Polidori *et al*., 2023; Youngsteadt *et al*., 2016). Across taxa, urban warming has been shown to alter phenology, abundance, and species turnover, and global analyses suggest that thermal stress should intensify toward lower latitudes where organisms operate closer to upper thermal limits (Deutsch *et al*., 2008; Diamond *et al*., 2012; Sunday *et al*., 2014). Based on these patterns, we predicted that urban heat would be a primary driver of species richness declines in Atlanta. However, surface temperature was not a significant predictor of native richness or composition. We cannot rule out temperature effects entirely, as work from Raleigh, North Carolina (USA), suggests urban heat can structure southeastern ant communities (Penick *et al*., 2025), but in our system, abiotic gradients were overshadowed by invasion pressure. Outside of temperature, species richness declined with increasing leaf litter cover and depth, factors typically associated with higher ant diversity under non-invaded conditions (e.g., Lassau & Hochuli, 2004; Queiroz *et al*., 2013; Silva *et al*., 2011). This counterintuitive relationship suggests that abiotic habitat features alone cannot explain the observed diversity pattern. Instead, leaf-litter-rich forests were also the habitats most heavily invaded by *B. chinensis*, indicating that biotic pressure may have overridden expected habitat effects.

Instead of abiotic stress, declines in richness and native abundance were strongly associated with *B. chinensis*, the most common ant in Atlanta based on species occurrence. Although first documented in the Atlanta metro area in 1932, *B. chinensis* recently expanded in range and abundance and only became prominent in Atlanta between 1991 and 2010 (Guénard *et al*., 2018). Unlike many urban invaders, *B. chinensis* readily persists in relatively undisturbed forests, including protected areas, such as Great Smoky Mountains National Park (Guénard & Dunn, 2010; Kanes *et al*., 2025). This ecological trait likely explains its disproportionate impact on forested parks in Atlanta. Based on patterns from New York City, where forested parks support ∼23% more species than manicured parks (Savage *et al*., 2015), we would have expected ∼49 species in Atlanta’s forested parks. Instead, we observed only 33 species, a ∼33% reduction. These losses are consistent with previous studies documenting ∼32% declines in richness and up to 70% reductions in native abundance in sites heavily invaded by *B. chinensis* (Guénard & Dunn, 2010). Patterns in native abundance followed similar trends in Atlanta, with an 80% decline in native ant abundance across the *B. chinensis* invasion gradient. These patterns suggest that *B. chinensis* exerts a substantial suppressive effect on native ant communities both in and outside of urban areas.

In contrast to *B. chinensis*, the two most economically damaging invasive ants in the southeastern United States, the red imported fire ant (*Solenopsis invicta*) and the Argentine ant (*Linepithema humile)*, had comparatively limited effects on native diversity in this study. Both species rank among the world’s 100 worst invasive alien species as determined by the International Union for Conservation of Nature (IUCN) (Lowe *et al*., 2000). Both species impose substantial economic costs due to damage and cost of management, with *S. invicta* alone estimated to have caused approximately $3 billion USD in damage in the United States since 1937 (Angulo *et al*., 2022). Yet neither species was particularly dominant across sites in Atlanta. We found that *S. invicta* was only the sixth most common ant species in Atlanta based on species occurrences, and was largely restricted to streetscapes and manicured parks; *L. humile* was rare based on species occurrences, but locally abundant in in the relatively few sites it was collected. While *S. invicta* and *L. humile* remain costly urban pests, particularly in highly disturbed habitats near structures, the forest-adapted *B. chinensis* exerted far stronger effects on native ant communities within remnant parks. These results underscore an important distinction between economic prominence and biodiversity impact, as relatively few resources have been deployed towards control of *B. chinensis* compared to these other invaders (Buczkowski, 2023).

Overall, our results suggest that the capacity of cities to function as protected habitats for insects may be reduced by invasive species, particularly at lower latitudes. If the patterns observed in Atlanta are representative, low-latitude cities may experience greater biodiversity declines associated with stronger invasion pressure. This interpretation is consistent with global analyses showing greater losses of ant diversity and increased divergence between urban and non-urban communities at lower latitudes (Perez *et al*., 2022), patterns attributed to higher dominance of non-native species, though not directly tested. In Atlanta, relatively intact forested parks did not provide refuge from invasion. This suggests that habitat protection alone may be insufficient to conserve native biodiversity in warmer regions where invasion pressure is high. Notably, Atlanta remains within a temperate region, underscoring the need for comparative studies in lower-latitude and tropical cities, where these dynamics may be even more pronounced and where the bioark hypothesis remains largely untested.

These findings have practical implications for urban conservation. Cities are major hubs for species introductions (Francis & Chadwick, 2015), and future urban expansion is projected to occur disproportionately in biodiverse regions (Potgieter *et al*., 2024; Seto *et al*., 2012). Yet most urban invasive ant management focuses on nuisance control around structures, where chemical baits are the primary treatment strategy (Dimarco *et al*., 2017). While chemical controls can be effective, they may generate non-target effects (Amaral *et al*., 2024; Hoffmann *et al*., 2023). Adjusting bait grain size, placement, timing, and formulation can further reduce collateral effects on native ants (Hoffmann *et al*., 2023; Vogt *et al*., 2005). Species-specific approaches may be particularly important for managing *B. chinensis*. Buczkowski (2016) demonstrated that termite-based “Trojan horse” baiting can selectively suppress *B. chinensis* with reduced non-target effects, although scaling such approaches may be challenging. More recently, Buczkowski (2023) showed that incorporating termite cuticular extracts into baits improves acceptance by *B. chinensis*, potentially increasing treatment efficacy. Seasonal timing may also matter, as bait acceptance by *B. chinensis* peaks between July and September (Corsetti, 2024). However, additional investment in effective control strategies for *B. chinensis* is needed to develop scalable methods at local eradication.

More broadly, our findings indicate that invasive insects can exert strong pressure even within relatively protected park habitats. Urban forests are often fragmented and subject to edge effects that facilitate invasion (Holway, 2005). Indeed, *B. chinensis* appears to perform particularly well in fragmented forests (Campbell *et al*., 2019). Cities may therefore act as incubators for forest-adapted invaders, not merely hubs for disturbance-associated species. The recent establishment of the ManhattAnt, *Lasius emarginatus* (Olivier, 1792), in New York City illustrates this broader pattern, as they were also first detected in urban parks before spreading into highly urban areas (Kennett *et al*., 2024). Cities are thus simultaneously potential bioarks and frontlines of biological invasion (Lin *et al*., 2025). In low-latitude regions where biodiversity is highest, preserving insect diversity will require not only protecting remnant habitats but actively managing invasive species within them. Without coordinated invasion control, remnant urban forests may fail to serve as refuges where they are most needed. Whether cities ultimately function as bioarks or sources of biodiversity loss will depend on our ability to manage invasion within these protected habitats.

## Acknowledgements

We thank Jenna Gaudet, Archel Sutton, Zachary Peagler, Layne Buttram, Rebecca Senft, and Amjad Alkawam for assistance with ant sampling in the field. We also appreciate Arthur Appel and Dan Warner for their valuable input on study design and presentation. Permission to sample ants in Atlanta parks was granted by the Atlanta Department of Parks and Recreation’s Bureau of Parks and Recreation.

## Data availability statement

All data and code required to recreate the analyses in this study are archived on Zenodo. DOI: https://doi.org/10.5281/zenodo.19568240

## Ethics approval statement

Ethical approval was not required for this study.

## References

1. Amaral, K.D., Marinho, C.G.S. & Della Lucia, T.M.C. (2024) Non-target ants and bioinsecticides: A short review. Current Opinion in Environmental Science & Health, 42, 100586.

2. Angulo, E., Hoffmann, B.D., Ballesteros-Mejia, L., Taheri, A., Balzani, P., Bang, A., et al. (2022) Economic costs of invasive alien ants worldwide. Biological Invasions, 24, 2041–2060.

3. Aronson, M.F.J., La Sorte, F.A., Nilon, C.H., Katti, M., Goddard, M.A., Lepczyk, C.A., et al. (2014) A global analysis of the impacts of urbanization on bird and plant diversity reveals key anthropogenic drivers. Proceedings of the Royal Society B: Biological Sciences, 281, 20133330.

4. Baldock, K.C.R., Goddard, M.A., Hicks, D.M., Kunin, W.E., Mitschunas, N., Osgathorpe, L.M., et al. (2015) Where is the UK’s pollinator biodiversity? The importance of urban areas for flower-visiting insects. Proceedings of the Royal Society B: Biological Sciences, 282, 20142849.

5. Brassard, F., Leong, C.-M., Chan, H.-H. & Guénard, B. (2021) High diversity in urban areas: how comprehensive sampling reveals high ant species richness within one of the most urbanized regions of the world. Diversity, 13, 358.

6. Buczkowski, G. (2016) The Trojan horse approach for managing invasive ants: a study with Asian needle ants, *Pachycondyla chinensis*. Biological Invasions, 18, 507–515.

7. Buczkowski, G. (2023) Termite cuticular extracts improve acceptance of bait for controlling invasive Asian needle ants, *Brachyponera chinensis*. Pest Management Science, 79, 4004–4010.

8. Cabon, V., Quénol, H., Dubreuil, V., Ridel, A. & Bergerot, B. (2024) Urban heat island and reduced habitat complexity explain spider community composition by excluding large and heat-sensitive species. Land, 13, 83.

9. Campbell, J.W., Grodsky, S.M., Halbritter, D.A., Vigueira, P.A., Vigueira, C.C., Keller, O., et al. (2019) Asian needle ant (*Brachyponera chinensis*) and woodland ant responses to repeated applications of fuel reduction methods. Ecosphere, 10, e02547.

10. Carpintero, S. & Reyes-López, J. (2008) The role of competitive dominance in the invasive ability of the Argentine ant (*Linepithema humile*). Biological Invasions, 10, 25–35.

11. Corsetti, K.A. (2024) *Bait acceptance and seasonal activity of the Asian needle ant*, Brachyponera *(=*Pachycondyla) chinensis *(Emery), an emerging medically important species, in central Georgia* (M.S.).

12. Deutsch, C.A., Tewksbury, J.J., Huey, R.B., Sheldon, K.S., Ghalambor, C.K., Haak, D.C., et al. (2008) Impacts of climate warming on terrestrial ectotherms across latitude. Proceedings of the National Academy of Sciences, 105, 6668–6672.

13. Dewitz, J. (2023) National Land Cover Database (NLCD) 2021 Products.

14. Diamond, S.E., Bellino, G. & Deme, G.G. (2023) Urban insect bioarks of the 21st century. Current Opinion in Insect Science, 57, 101028.

15. Diamond, S.E., Nichols, L.M., McCoy, N., Hirsch, C., Pelini, S.L., Sanders, N.J., et al. (2012) A physiological trait-based approach to predicting the responses of species to experimental climate warming. Ecology, 93, 2305–2312.

16. Dietzel, A., Moretti, M., Perrelet, K. & Cook, L.M. (2025) Urban heat exacerbates climatic risks to urban biodiversity. npj Urban Sustainability, 6, 4.

17. Dimarco, R.D., Masciocchi, M. & Corley, J.C. (2017) Managing nuisance social insects in urban environments: an overview. International Journal of Pest Management, 63, 251–265.

18. Fenoglio, M.S., Rossetti, M.R. & Videla, M. (2020) Negative effects of urbanization on terrestrial arthropod communities: A meta-analysis. Global Ecology and Biogeography, 29, 1412–1429.

19. Francis, R.A. & Chadwick, M.A. (2015) Urban invasions: non-native and invasive species in cities. Geography, 100, 144–151.

20. Gochnour, B.M., Suiter, D.R. & Booher, D. (2019) Ant (hymenoptera: formicidae) fauna of the marine port of Savannah, Garden City, Georgia (USA). Journal of Entomological Science, 54, 417–429.

21. Gotelli, N.J., Ellison, A.M., Dunn, R.R. & Sanders, N.J. (2011) Counting ants (Hymenoptera: Formicidae): Biodiversity sampling and statistical analysis for myrmecologists. College of Arts and Sciences Faculty Publications.

22. Griffith, J., Sunday, J.M. & Hargreaves, A.L. (2025) Urbanization weakens the latitudinal diversity gradient in birds.

23. Gruber, M.A.M., Janssen-May, S., Santoro, D., Cooling, M. & Wylie, R. (2021) Predicting socio-economic and biodiversity impacts of invasive species: Red Imported Fire Ant in the developing western Pacific. Ecological Management & Restoration, 22, 89–99.

24. Guénard, B. & Dunn, R.R. (2010) A new (old), invasive ant in the hardwood forests of eastern north america and its potentially widespread impacts. PLOS ONE, 5, e11614.

25. Guénard, B., Wetterer, J.K. & MacGown, J.A. (2018) Global and temporal spread of a taxonomically challenging invasive ant, *Brachyponera chinensis* (Hymenoptera: Formicidae). Florida Entomologist, 101, 649–656.

26. Guo, Q., Cade, B.S., Dawson, W., Essl, F., Kreft, H., Pergl, J., et al. (2021) Latitudinal patterns of alien plant invasions. Journal of Biogeography, 48, 253–262.

27. Hoffmann, B.D., Pettit, M., Antonio, J., Chassain, J., Ferrieu, E., Gutierrez, A., et al. (2023) Efficacy, non-target impacts, and other considerations of unregistered fipronil-laced baits being used in multiple invasive ant eradication programs. Management of Biological Invasions, 14, 437–457.

28. Holway, D.A. (2005) Edge effects of an invasive species across a natural ecological boundary. Biological Conservation, 121, 561–567.

29. Holway, D.A., Lach, L., Suarez, A.V., Tsutsui, N.D. & Case, T.J. (2002) The causes and consequences of ant invasions. Annual Review of Ecology, Evolution, and Systematics, 33, 181–233.

30. Kanes, D., Malagon, D., Camper, B., Hewitt, A., Dunn, S., Purcell, E., et al. (2025) Species distribution models reveal varying degrees of refugia from the invasive Asian needle ant for native ants versus ant-plant seed dispersal mutualisms. Ecology and Evolution, 15, e70750.

31. Kennett, S.M., Seifert, B., Dunn, R.R., Pierson, T.W. & Penick, C.A. (2024) The ManhattAnt: identification, distribution, and colony structure of a new pest in New York City, *Lasius emarginatus*. Biological Invasions, 26, 2759–2772.

32. Lassau, S.A. & Hochuli, D.F. (2004) Effects of habitat complexity on ant assemblages. Ecography, 27, 157–164.

33. Leveau, L.M., Leveau, C.M., Villegas, M., Cursach, J.A. & Suazo, C.G. (2017) Bird communities along urbanization gradients: a comparative analysis among three neotropical cities. Ornitología Neotropical, 28, 77–87.

34. Lin, W.-J., Hsu, P.-W., Vargo, E.L. & Yang, C.-C.S. (2025) Microbial, genetic, and urban drivers of ant invasions. Current Opinion in Insect Science, 72, 101417.

35. Lowe, S., Browne, M., Boudjelas, S. & De Poorter, M. (2000) 100 of the world’s worst invasive alien species: a selection from the global invasive species database. Invasive Species Specialist Group Auckland.

36. MacGown, J. (2024) Ants (Formicidae) of the Southeastern United States. https://mississippientomologicalmuseum.org.msstate.edu/Researchtaxapages/Formicidaepages/Identification.Keys.htm [accessed on 2024].

37. Mignanelli, M. (2021) Percentage Cover.

38. Oksanen, J., Simpson, G.L., Blanchet, F.G., Kindt, R., Legendre, P., Minchin, P.R., et al. (2025) vegan: Community Ecology Package.

39. Penick, C., Peagler, Z., Buttram, L., Dunn, R., Frank, S. & Youngsteadt, E. (2025) Urban heat and latitude: contrasting effects on ant diversity across cities. Urban Ecosystems, 28.

40. Penick, C.A. (2021) Urban social insects. In: Encyclopedia of social insects. Springer, pp. 983–988.

41. Perez, A., Chick, L., Menke, S., Lessard, J.-P., Sanders, N., Del Toro, I., et al. (2022) Urbanisation dampens the latitude-diversity cline in ants. Insect Conservation and Diversity, 15, 763–771.

42. Polidori, C., Ferrari, A., Ronchetti, F., Tommasi, N. & Nalini, E. (2023) Warming up through buildings and roads: what we know and should know about the urban heat island effect on bees. Frontiers in Bee Science, 1.

43. Potgieter, L.J., Li, D., Baiser, B., Kühn, I., Aronson, M.F.J., Carboni, M., et al. (2024) Cities shape the diversity and spread of nonnative species. Annual Review of Ecology, Evolution, and Systematics, 55, 157–180.

44. Queiroz, A.C.M. de, Ribas, C.R. & França, F.M. (2013) Microhabitat characteristics that regulate ant richness patterns: the importance of leaf litter for epigaeic ants. Sociobiology, 60, 367–373.

45. R Core Team. (2025) R: a language and environment for statistical computing. R Foundation for Statistical Computing.

46. Rajesh, T.P., Manoj, K., Prashanth Ballullaya, U., Shibil, V.K., Asha, G., Varma, S., et al. (2022) Urban tropical forest islets as hotspots of ants in general and invasive ants in particular. Scientific Reports, 12, 12003.

47. Sanllorente, O., Blanco-Urdillo, E., Sánchez-Tójar, A. & Diego Ibáñez-Álamo, J. (2025) A systematic review and meta-analysis on urban arthropod diversity. Insect Conservation and Diversity, 18, 447–464.

48. Sattler, T., Obrist, M.K., Duelli, P. & Moretti, M. (2011) Urban arthropod communities: Added value or just a blend of surrounding biodiversity? Landscape and Urban Planning, 103, 347.

49. Savage, A.M., Hackett, B., Guénard, B., Youngsteadt, E.K. & Dunn, R.R. (2015) Fine-scale heterogeneity across Manhattan’s urban habitat mosaic is associated with variation in ant composition and richness. Insect Conservation and Diversity, 8, 216–228.

50. Sears, M., Angilletta, M., Wilson, R., Niehaus, A., Ribeiro, P. & Navas, C. (2007) Urban physiology: city ants possess high heat tolerance. PLOS ONE.

51. Senator, S.A. & Rozenberg, A.G. (2017) Assessment of economic and environmental impact of invasive plant species. Biology Bulletin Reviews, 7, 273–278.

52. Seto, K.C., Güneralp, B. & Hutyra, L.R. (2012) Global forecasts of urban expansion to 2030 and direct impacts on biodiversity and carbon pools. Proceedings of the National Academy of Sciences, 109, 16083–16088.

53. Silva, P.S.D., Bieber, A.G.D., Corrêa, M.M. & Leal, I.R. (2011) Do leaf-litter attributes affect the richness of leaf-litter ants? Neotropical Entomology, 40, 542–547.

54. Sunday, J.M., Bates, A.E., Kearney, M.R., Colwell, R.K., Dulvy, N.K., Longino, J.T., et al. (2014) Thermal-safety margins and the necessity of thermoregulatory behavior across latitude and elevation. Proceedings of the National Academy of Sciences, 111, 5610–5615.

55. Terando, A.J., Costanza, J., Belyea, C., Dunn, R.R., McKerrow, A. & Collazo, J.A. (2014) The southern megalopolis: using the past to predict the future of urban sprawl in the southeast U.S. PLOS ONE, 9, e102261.

56. U.S. Geological Survey. (2021) Landsat Collectoin 2 Level-2 Science Products ( No. 2021–3055). Fact Sheet.

57. Vogt, J.T., Reed, J.T. & Brown, R.L. (2005) Timing bait applications for control of imported fire ants (Hymenoptera: Formicidae) in Mississippi: Efficacy and effects on non-target ants. International journal of pest management, 51, 121–130.

58. Wallace, A.R. (1878) Tropical nature, and other essays. Macmillan and Company.

59. Warren, R.J., McMillan, A., King, J.R., Chick, L. & Bradford, M.A. (2015) Forest invader replaces predation but not dispersal services by a keystone species. Biological Invasions, 17, 3153–3162.

60. Youngsteadt, E., Ernst, A.F., Dunn, R.R. & Frank, S.D. (2016) Responses of arthropod populations to warming depend on latitude: evidence from urban heat islands. Global Change Biology, 23, 1436–1447.

