## Supplement for "Invasive species undermine the bioark hypothesis in a low-latitude urban biodiversity hotspot"

### Supplementary material

**Table S1.** Site location and sampling methods

| Site | Habitat type | Latitude | Longitude | Sampling method(s) |
|---|---|---|---|---|
| S01 | Forested park | 33.7875 | -84.371944 | Hand, bait, winkler |
| S02 | Forested park | 33.791111 | -84.371944 | Hand, bait, winkler |
| S03 | Forested park | 33.790414 | -84.382421 | Hand, bait, winkler, pitfall |
| S04 | Manicured park | 33.78857 | -84.380784 | Hand, bait, winkler |
| S05 | Streetscape | 33.788456 | -84.383533 | Hand, bait, winkler |
| S06 | Streetscape | 33.794966 | -84.387935 | Hand, bait, winkler, pitfall |
| S07 | Manicured park | 33.7825 | -84.375556 | Hand, bait, winkler |
| S08 | Manicured park | 33.7825 | -84.375556 | Hand, bait, winkler |
| S09 | Forested park | 33.782787 | -84.375558 | Hand, bait, winkler, pitfall |
| S11 | Manicured park | 33.769551 | -84.352222 | Hand, bait, winkler, pitfall |
| S12 | Forested park | 33.772462 | -84.336777 | Hand, bait, winkler |
| S13 | Forested park | 33.793338 | -84.363592 | Hand, bait, winkler |
| S14 | Forested park | 33.785469 | -84.361223 | Hand, bait, winkler, pitfall |
| S15 | Forested park | 33.7858 | -84.363114 | Hand, bait, winkler, pitfall |
| S16 | Forested park | 33.795475 | -84.374626 | Hand, bait, winkler, pitfall |
| S17 | Streetscape | 33.7925 | -84.386944 | Hand, bait, winkler, pitfall |
| S18 | Streetscape | 33.790278 | -84.387222 | Hand, bait, winkler, pitfall |
| S19 | Forested park | 33.792438 | -84.377936 | Hand, bait, winkler |
| S20 | Streetscape | 33.787796 | -84.387486 | Hand, bait, winkler |
| S21 | Streetscape | 33.754937 | -84.388715 | Hand, bait, winkler |
| S22 | Manicured park | 33.770658 | -84.347666 | Hand, bait, winkler, pitfall |
| S23 | Manicured park | 33.760184 | -84.394471 | Hand, bait, winkler, pitfall |
| S24 | Manicured park | 33.760184 | -84.394471 | Hand, bait, winkler, pitfall |
| S25 | Streetscape | 33.782778 | -84.384167 | Hand, bait, winkler |
| S26 | Streetscape | 33.782302 | -84.383878 | Hand, bait, winkler |
| S27 | Manicured park | 33.761365 | -84.394063 | Hand, bait, winkler |
| S28 | Streetscape | 33.755661 | -84.388431 | Hand, bait, winkler |
| S30 | Streetscape | 33.758475 | -84.402273 | Hand, bait, winkler, pitfall |
| S31 | Streetscape | 33.784041 | -84.382932 | Hand, bait, winkler |
| S32 | Streetscape | 33.767653 | -84.354766 | Hand, bait, winkler, pitfall |

| Site | Habitat type | Latitude | Longitude | Sampling method(s) |
|---|---|---|---|---|
| S33 | Streetscape | 33.759582 | -84.389542 | Hand, bait, winkler, pitfall |
| S35 | Manicured park | 33.769029 | -84.358211 | Hand, bait, winkler, pitfall |
| S37 | Forested park | 33.758045 | -84.357692 | Hand, bait, winkler, pitfall |
| S38 | Forested park | 33.757785 | -84.357449 | Hand, bait, winkler, pitfall |
| S39 | Manicured park | 33.784492 | -84.378273 | Hand, bait, winkler |
| S40 | Manicured park | 33.769838 | -84.346813 | Hand, bait, winkler |
| S41 | Forested park | 33.771101 | -84.343492 | Hand, bait, winkler |
| S42 | Manicured park | 33.758305 | -84.35788 | Hand, bait, winkler, pitfall |
| S43 | Forested park | 33.793736 | -84.3763 | Hand, bait, winkler, pitfall |
| S44 | Forested park | 33.793188 | -84.376863 | Hand, bait, winkler |
| S45 | Forested park | 33.768662 | -84.356275 | Hand, bait, winkler |
| S46 | Forested park | 33.7660605 | -84.3376181 | Hand, bait, winkler |
| S47 | Manicured park | 33.7660605 | -84.3376181 | Hand, bait, winkler |
| S48 | Manicured park | 33.7366051 | -84.3722826 | Hand, bait, winkler |
| S49 | Manicured park | 33.7393781 | -84.3734571 | Hand, bait, winkler |
| S51 | Streetscape | 33.755519 | -84.359384 | Hand, bait, winkler, pitfall |
| S52 | Streetscape | 33.758307 | -84.393585 | Hand, bait, winkler |
| S101 | Streetscape | 33.774225 | -84.416938 | Pitfall |
| S102 | Streetscape | 33.781395 | -84.399355 | Pitfall |
| S103 | Manicured park | 33.770191 | -84.376930 | Pitfall |
| S105 | Manicured park | 33.766500 | -84.337191 | Pitfall |
| S106 | Forested park | 33.738082 | -84.371237 | Pitfall |
| S107 | Manicured park | 33.736457 | -84.369385 | Pitfall |
| S108 | Forested park | 33.790225 | -84.368503 | Pitfall |
| S109 | Forested park | 33.790335 | -84.366770 | Pitfall |
| S110 | Manicured park | 33.789940 | -84.368322 | Pitfall |
| S111 | Manicured park | 33.789886 | -84.366507 | Pitfall |
| S112 | Forested park | 33.793308 | -84.363614 | Pitfall |
| S113 | Streetscape | 33.782159 | -84.383785 | Pitfall |
| S114 | Forested park | 33.773726 | -84.338143 | Pitfall |

**Table S2.** Occurrence data for all species found during hand collection, winkler extraction and baiting. There were 18 total streetscapes, 20 total manicured parks, and 21 total forested parks for a total of 59 total sites. Non-native species are indicated by an asterisk (*).

| Species | Streetscape | Manicured Park | Forested Park | Total Sites Found |
|---|---|---|---|---|
| *Aphaenogaster flemingi* Smith | 1 | 0 | 0 | 1 |
| *Aphaenogaster fulva* Roger | 0 | 1 | 1 | 2 |
| *Aphaenogaster rudis complex* Wesson & Wesson | 0 | 1 | 0 | 1 |
| *Brachymyrmex depilis* Emery | 0 | 1 | 1 | 2 |
| *Brachymyrmex obscurior* Forel * | 1 | 3 | 1 | 5 |
| *Brachymyrmex patagonicus* Mayr * | 10 | 6 | 1 | 17 |
| *Brachymyrmex sp.02* | 1 | 1 | 2 | 4 |
| *Brachyponera chinensis* * (Emery) | 12 | 17 | 21 | 50 |
| *Camponotus americanus* Mayr | 0 | 0 | 2 | 2 |
| *Camponotus castaneus* Latreille | 5 | 6 | 12 | 23 |
| *Camponotus chromaiodes* Bolton | 0 | 0 | 2 | 2 |
| *Camponotus nearcticus* Emery | 1 | 1 | 0 | 2 |
| *Camponotus pennsylvanicus* (De Geer) | 3 | 12 | 16 | 31 |
| *Camponotus snellingi* Bolton | 3 | 0 | 5 | 8 |
| *Crematogaster ashmeadi* Mayr | 5 | 8 | 6 | 19 |
| *Crematogaster lineolata* (Say) | 0 | 1 | 0 | 1 |
| *Crematogaster minutissima* Mayr | 1 | 0 | 1 | 2 |
| *Crematogaster missouriensis* Emery | 1 | 2 | 0 | 3 |
| *Crematogaster vermiculata* Emery | 1 | 0 | 1 | 2 |
| *Forelius mccooki* (McCook) | 0 | 1 | 0 | 1 |
| *Formica pallidefulva* Latreille | 4 | 10 | 12 | 26 |
| *Formica rubicunda* Emery | 0 | 2 | 3 | 5 |
| *Formica subsericea* Say | 1 | 3 | 3 | 7 |
| *Hypoponera punctatissima* (Roger) * | 0 | 1 | 0 | 1 |
| *Lasius americanus* Emery | 10 | 14 | 8 | 32 |
| *Lasius aphidicola* (Walsh) | 0 | 0 | 1 | 1 |
| *Lasius nearcticus* Wheeler | 0 | 1 | 0 | 1 |
| *Lasius neoniger* Emery | 5 | 16 | 12 | 33 |
| *Linepithema humile* (Mayr) * | 5 | 2 | 0 | 7 |
| *Monomorium carbonarium* (Smith) | 10 | 4 | 3 | 17 |

| Species | Streetscape | Manicured Park | Forested Park | Total Sites Found |
|---|---|---|---|---|
| *Monomorium viridum* Brown | 1 | 1 | 0 | 2 |
| *Myrmecina americana* Emery | 1 | 0 | 0 | 1 |
| *Nylanderia faisonensis* (Forel) | 0 | 4 | 0 | 4 |
| *Nylanderia wojciki* (Trager) | 15 | 14 | 7 | 36 |
| *Pheidole bicarinata* Mayr | 4 | 2 | 2 | 8 |
| *Pheidole dentata* Mayr | 1 | 3 | 0 | 4 |
| *Pheidole dentigula* Smith | 0 | 0 | 1 | 1 |
| *Pheidole megacephala* (Fabricius) * | 0 | 1 | 0 | 1 |
| *Pheidole morrisii* Forel | 0 | 1 | 0 | 1 |
| *Pheidole tysoni* Forel | 7 | 12 | 8 | 27 |
| *Prenolepis imparis* (Say) | 0 | 0 | 1 | 1 |
| *Pseudomyrmex ejectus* (Smith) | 0 | 1 | 0 | 1 |
| *Solenopsis invicta* Buren * | 13 | 8 | 5 | 26 |
| *Solenopsis molesta* (Say) | 12 | 7 | 5 | 24 |
| *Stigmatomma pallipes* (Haldeman) | 1 | 0 | 0 | 1 |
| *Strumigenys archboldi* (Deyrup & Cover) | 0 | 0 | 1 | 1 |
| Strumigenys hexamera (Brown) * | 1 | 0 | 0 | 1 |
| *Strumigenys louisianae* Roger | 1 | 2 | 3 | 6 |
| *Strumigenys membranifera* Emery * | 3 | 2 | 1 | 6 |
| *Strumigenys* ornata Mayr | 0 | 0 | 1 | 1 |
| *Strumigenys silvestrii* (Emery) * | 0 | 1 | 0 | 1 |
| *Tapinoma sessile* (Say) | 7 | 9 | 6 | 22 |
| *Temnothorax schaumii* (Roger) | 0 | 2 | 0 | 2 |
| *Trachymyrmex septentrionalis* (McCook) | 0 | 1 | 0 | 1 |

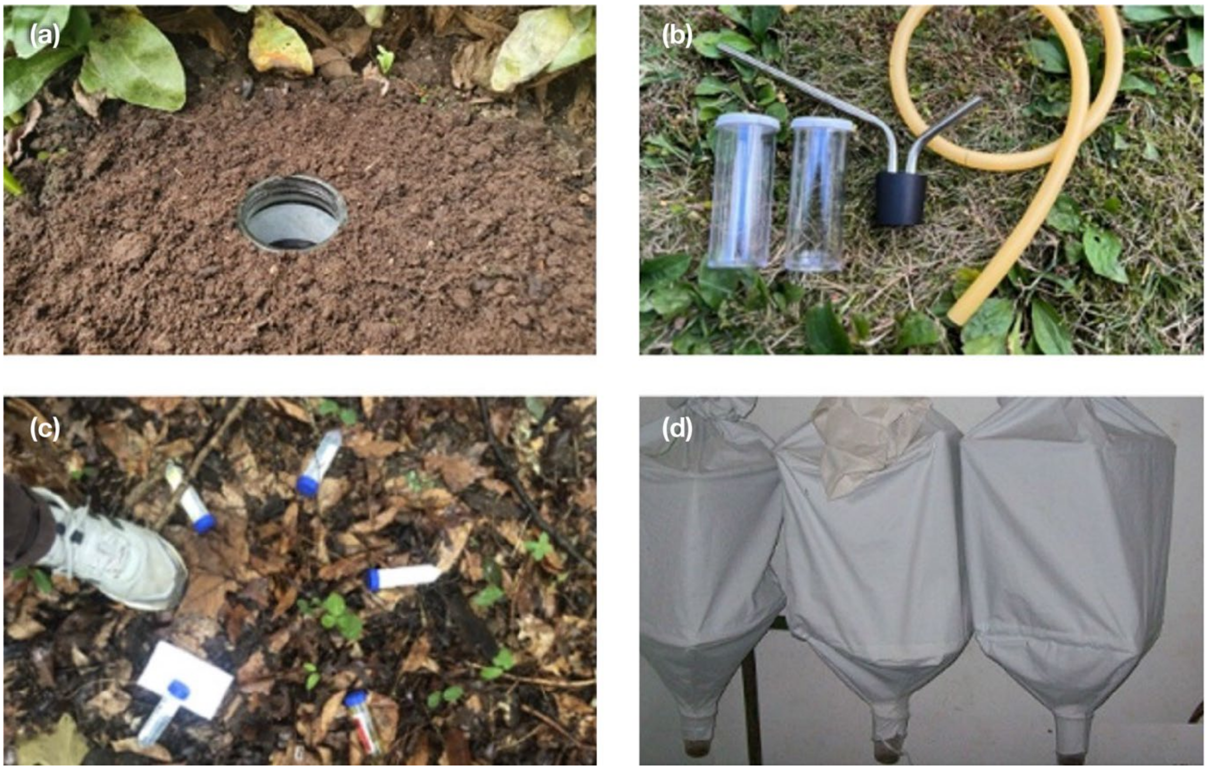


**Fig. S1** Sampling methods. **(a)** Pitfall trap filled with soapy water, **(b)** aspirator used for hand sampling, **(c)** liquid baits, and (**d)** Winkler bags used to extract ants from leaf litter samples.

**Figure S2.** Relationships between ant species richness and six abiotic and environmental variables across all sites. **(a)** Canopy cover, **(b)** impervious surface cover, **(c)** leaf litter cover, **(d)** leaf litter depth, **(e)** maximum surface temperature, and **(f)** vegetation complexity (Shannon index, H′). Each point represents one site. Linear regression lines with 95% confidence intervals are shown for significant relationships ($p < 0.05$). $p$-values are from linear models.

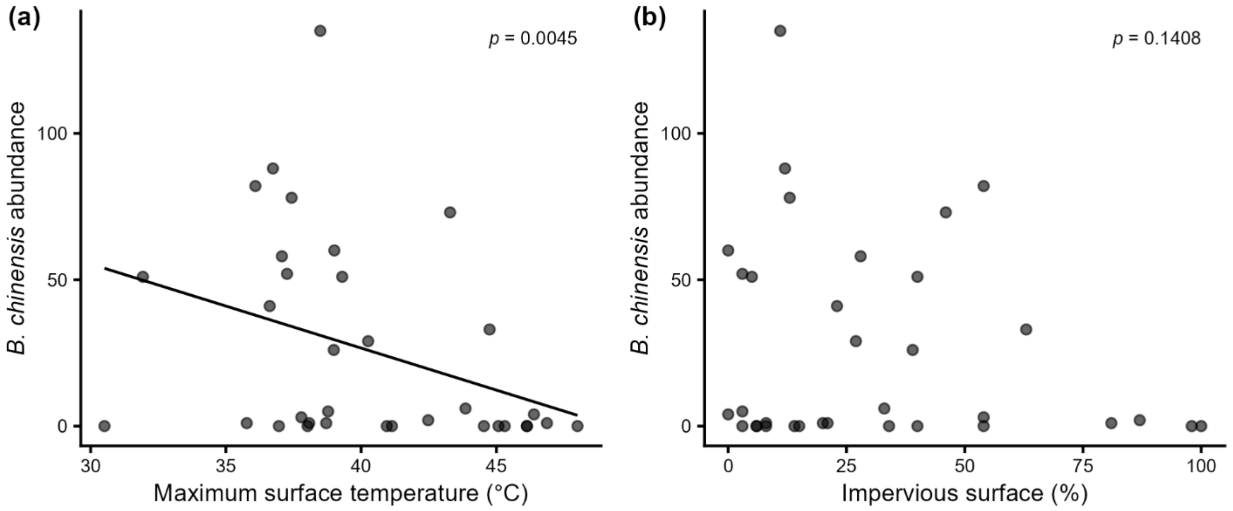


**Fig. S3** Relationship between *Brachyponera chinensis* pitfall trap abundance and (**a**) maximum surface temperature (°C) and (**b**) impervious surface cover (%) across 34 sites. Both relationships were modelled using negative binomial generalised linear models to account for overdispersion in count data. A fitted linear trend line is shown where the relationship was statistically significant ($p < 0.05$).
